# Proteome-wide multipoint internal calibration curves for mass spectrometry-based quantitative proteomics

**DOI:** 10.1101/2025.02.24.639845

**Authors:** Cristina Chiva, Zahra Elhamraoui, Julia Morales-Sanfrutos, Olga Pastor, Eduard Sabidó

## Abstract

Mass spectrometry (MS)-based proteomics is known for its high accuracy in quantifying peptides and proteins using various calibration strategies, including internal and external calibration curves. While external multi-point calibration curves are created from serial dilutions, they often fail to account for sample-specific matrix effects. In contrast, internal calibration curves account for sample matrix but face scalability and cost challenges for whole proteome analyses. In this manuscript we present a novel TMT-based multipoint internal calibration curve strategy, referred to as TMTCal, which enables the generation of internal calibration curves for all peptides identified within a proteome within a single experiment. We applied this strategy to human ovarian cancer cells to evaluate the linear quantitative responses of all the identified peptides and reveal the significant proteome changes associated with cisplatin treatment.

---

Mass spectrometry (MS)-based proteomics is well-known for its ability to quantify peptides and proteins with high accuracy and precision.^1,2^ In quantitative targeted proteomics experiments, various calibration strategies are employed, including internal and external calibration curves, to establish the relationship between peptide signal and peptide concentration.^3–6^ External multi-point calibration curves are created through serial dilutions of a standard containing known concentrations of the peptide of interest. While these curves establish the fundamental relationship between instrument response and analyte concentration, they have limitations in accounting for sample-specific matrix effects, as the range of linearity is not directly determined within the sample. This issue is particularly significant when dealing with patient cohorts, where variations in sample matrices can affect quantification accuracy. To address these limitations, internal calibration curves have been proposed, using either matrix-matched approaches^7^ or by spiking isotopically labeled standards directly into the samples.^8–10^ These internal isotopically labeled standards are often used in the single-point internal calibration mode, in which a known concentration of the standard is added to determine the response factor, assuming a linear relationship through zero. While this method is quick and resource-efficient, it does not fully capture the complexity of the response curve. A few years ago we developed an isotopologue multipoint calibration strategy (ImCal) that uses a mixture of isotopically labeled peptides at different concentrations to establish a multipoint internal calibration curve.^11^ This method ensured precise and accurate quantification of specific peptides in targeted proteomics applications.^11–13^ However, the use of multiple isotopologue peptides increases the cost of each targeted assay, and its application to entire proteomes remains limited due to the challenges of scaling, making it suitable only for clinical projects in which a small number of peptides are measured across a large number of samples.

In this study, we expanded the ImCal concept to the entire proteome by developing a tandem mass tag (TMT)- based multipoint internal calibration curve strategy for all peptides identified within a proteome. This approach, which we refer to as TMTCal, is based on TMT-labeled serial dilutions of total protein extract that are used to generate internal calibration curves together with the samples of interest within a single experiment, thus extending the advantages of ImCal quantitation to whole-proteome analyses.

To achieve this, human ovarian cancer cells (SK-OV-3) were cultured in triplicate, with and without cisplatin (25 µM), and the resulting proteomes were digested with endoproteinase LysC and trypsin. Simultaneously, a pooled sample was prepared to generate serial dilutions for the internal multipoint calibration curve (**Figure 1A** and **1B**, and **Table 1**). Tandem mass tags (TMT-11) were employed to label all the samples and serial dilutions, which were combined to form a single multiplexed TMT experiment. After basic pH reversed-phase fractionation, the TMT samples were analyzed using a 90-minute gradient on an Orbitrap Eclipse mass spectrometer, using an acquisition method with real-time search and synchronous precursor selection MS3 analysis (RTS-SPS-MS3).^14,15^ The acquired data were processed using Proteome Discoverer (v2.4) and all the mass spectrometry proteomics data have been deposited to the ProteomeXchange Consortium via the PRIDE partner repository^16^ with the dataset identifier PXD059628 (see detailed methods in **Supporting Information**).

**Figure 1:**
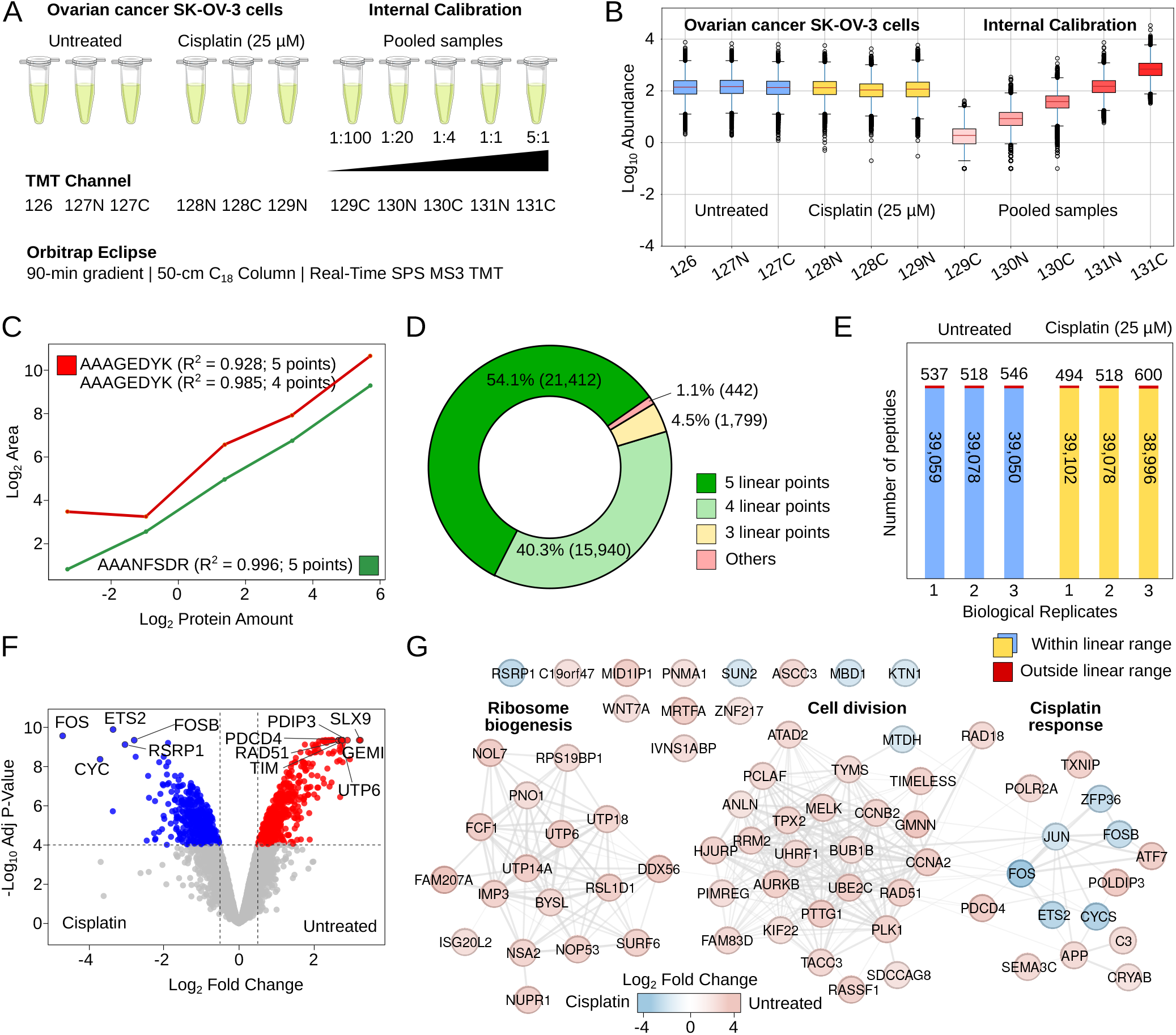
A) Human ovarian cancer cells (SK-OV-3) were cultured in triplicate with and without cis-platinum treatment (25 µM) and subsequently digested with trypsin. An internal multipoint calibration curve was generated using a pooled sample with six different dilutions ranging from 5:1 to 1:100. B) Experimental log-transformed peptide abundance values corresponding to the three biological replicates of human ovarian cancer cells (SK-OV-3) cultured with and without cis-platinum, and to the serial dilutions of the internal calibration curve. C) Linear regression fit for peptides [K].AAANFSDR.[S] (green) with 5 linear points (R^2^ > 0.99), and [K].AAAGEDYK.[A] (red) with four linear points (R^2^ > 0.98). D) Donut chart illustrating the classification of peptide linearity based on the count of linear points with R^2^ > 0.95. A linear fit was applied to all valid quantitative points for each peptide, and R^2^ was calculated. If R^2^ was below 0.95, the lowest concentration value was iteratively removed until R^2^ exceeded 0.95. The remaining quantitative values represent the number of linear points for that peptide. E) Number of endogenous peptides from the ovarian cancer cell lines that lay within the linear range established with the internal multipoint calibration curves. F) Volcano plot with proteins with significant abundance changes in the human ovarian cancer cells (SK-OV-3) with and without cis-platinum treatment (25 µM). G) Functional protein-protein interaction networks (String-db) with the proteins exhibiting the most significant fold-changes in protein abundance in cis-platinum treated human ovarian cancer cells (SK-OV-3).

**Table 1.** Experimental design for samples, tandem mass tags, and protein amounts (µg) multiplexed in a single TMT batch.

| TMT channel | Sample | Amount (μg) |
| --- | --- | --- |
| 126 | SK-OV-3 Cells Untreated Replicate 1 | 25 |
| 127N | SK-OV-3 Cells Untreated Replicate 2 | 25 |
| 127C | SK-OV-3 Cells Untreated Replicate 3 | 25 |
| 128N | SK-OV-3 Cells Treated (25 μM cisplatin) Replicate 1 | 25 |
| 128C | SK-OV-3 Cells Treated (25 μM cisplatin) Replicate 2 | 25 |
| 129N | SK-OV-3 Cells Treated (25 μM cisplatin) Replicate 3 | 25 |
| 129C | Calibration curve pool | 0.25 |
| 130N | Calibration curve pool | 1.25 |
| 130C | Calibration curve pool | 6.25 |
| 131N | Calibration curve pool | 25 |
| 131C | Calibration curve pool | 125 |

Using the TMTCal approach, we first evaluated the linear behavior of the TMT-labeled internal calibration curves for 39,596 peptides with at least three valid quantitation values—out of a total of 46,641 identified peptides (**Supplementary Table S1** and **Supporting Information**). Based on the log-transformed MS3 signals obtained from the TMT-labeled calibration curves, we calculated the number of linear points for each peptide. Briefly, a linear fit was applied to all valid quantitative points for each peptide and R^2^ was calculated. If R^2^ was below 0.95, the lowest concentration value was iteratively removed until R^2^ exceeded 0.95. The remaining quantitative values represent the number of linear points for that peptide. With this approach, we observed that the majority of identified peptides exhibited an excellent linear quantitative response, with approximately 95% of them demonstrating complete or nearly complete linear regression curves, i.e. four or five linear points with R^2^>0.95 (**Figure 1C** and **1D**). Not only were most calibration curves complete, but when missing values did occur in peptides with at least one quantitation value, these missing values were predominantly found at the lowest values of the calibration curves (i.e. 89.7% of the missing points were found in the lowest concentration), thus aligning with the expectations for the data. The excellent linear quantitative response observed in this study stands in contrast to previous quantitative results reported at the MS1 and MS2 levels,^7,17,18^ and it is likely attributable to the enhanced interference removal achieved through the synchronous precursor selection (SPS) MS3 quantification strategy employed in this study.

After evaluating the linear behavior of the TMT-labeled dilution curves for all identified peptides, we aimed to assess the reproducibility of this quantitative behavior across different batches. To achieve this, we conducted two additional TMT-labeled experiments, following the same experimental design, sample preparation, data acquisition, and data analysis protocols as previously described. The analysis of the three TMT-calibrated replicate experiments resulted in the identification and quantification of 57,951 peptides, with approximately 70% (39,224 peptides) being identified and quantified in at least two out of the three batches (**Supplementary Figure S1**). This result highlights a common challenge in analyzing multiple TMT batches, where slight variations in the list of identified peptides reduce completeness across batches. Nevertheless, for the peptides identified in all batches, we evaluated whether their quantitative behavior remained consistent. We observed that approximately 90% (35,077) peptides exhibiting complete or nearly complete linear regression dilution curves, i.e. four or five linear points with R^2^ >0.95 in one batch, also demonstrated a complete or nearly complete linear quantitative behavior in the other batches in which they were identified. Some peptides exhibited four linear quantitative points in one batch, while showing five in another, and vice versa. In the batches where only four linear quantitative points are observed, the peptides consistently display lower intensity in the MS3 spectra compared to the same peptide in the other TMTCal batches. This observation could be attributed to the precise timing of the sampling events for fragmentation during chromatographic elution, to small differences in R^2^ values around 0.95, and to specific matrix effects from co-eluting peptides present in each of the three batches (**Supplementary Figure S2**).

Finally, we applied the TMTCal calibration curves to a model system of ovarian cancer (SK-OV-3 cell line) treated with and without cisplatin (25 µM) to quantify the proteome remodeling that occurs after treatment. First, we assessed whether the endogenous peptides fell within the linear range of quantification established by the internal multipoint calibration curves constructed for each peptide. We confirmed that the vast majority of peptides identified in the human ovarian cancer cells were indeed within the linear range of quantification (**Figure 1E**). This observation ensured accurate and precise relative quantification either through direct comparison of the MS3 reported ion intensities, or by adjusting them using the individualized regression lines from the multipoint calibration curves available for each peptide. As a direct consequence of this finding, the fold changes in protein abundances obtained from direct reporter ion signals comparisons were very similar to those derived from calibrated reporter ion signals (**Supplementary Figure S3**). Despite the calibrated approach did not yield a significant advantage in this experiment involving whole cell extracts at large quantities, it holds great potential for enhancing quantification results in more sensitivity-challenging scenarios, such as low-input and single-cell proteomics. Lastly, we conducted relative quantification statistical inference of all the endogenous identified proteins with MSstats (v4.12.1).^19–21^ This relative quantification allowed us to discern the significant changes in the proteome of the ovarian cancer cell lines following cisplatin treatment, which primarily affected proteins associated with the ribosome biogenesis, cell cycle, nucleic acid metabolic processes and, as expected, the cellular response to cisplatin (**Figure 1F and 1G, Supplementary Figure S4**, and **Supplementary Table ST3**).

In conclusion, in this study we developed a tandem mass tag (TMT)-based multipoint internal calibration curve strategy, termed TMTCal, which extends the benefits of internal calibration to whole-proteome analyses. By applying this approach to human ovarian cancer cells together with real-time synchronous precursor selection (SPS) MS3 quantification, we demonstrated that the majority of identified peptides exhibited excellent linear quantitative responses, ensuring accurate and precise quantification. Our findings highlight the potential of TMTCal to enhance quantification in challenging scenarios, such as low-input and single-cell proteomics, while also providing insights into proteome remodeling in response to cisplatin treatment.

## Acknowledgments

We acknowledge support of the Spanish Ministry of Science and Innovation through the Centro de Excelencia Severo Ochoa (CEX2020-001049-S grant funded by MCIN/AEI/10.13039/501100011033) and PID2020-115092GB-I00 funded by AEI/10.13039/501100011033, and the Generalitat de Catalunya through the CERCA programme and the Departament de Recerca i Universitats (2021-SGR2021-01225). We also thank MD Melissa Bradbury for her valuable advice on the experimental design of the conditions for cisplatin treatment. This result is part of a project that has received funding from the European Union’s Horizon 2020 research and innovation programme under the Marie Sklodowska-Curie grant agreement No 956148. The CRG/UPF Proteomics Unit is part of the Spanish Infrastructure for Omics Technologies (ICTS OmicsTech).

