## Supporting Information for "Proteome-wide multipoint internal calibration curves for mass spectrometry-based quantitative proteomics"

### Table of Contents

- Materials and Methods
- Supplementary Table 1
- Supplementary Table 2
- Supplementary Table 3
- Supplementary Figure 1
- Bibliography

### Materials and methods

Human ovarian cancer cells (SK-OV-3 from American Type Culture Collection, ATCC) were cultured in presence or absence of 25 µM cisplatin in triplicates. Protein extracts (600 µg) were reduced with TCEP (1.8 µmol, 37 ºC, 60 min), alkylated in the dark with iodoacetamide (3.6 µmol, 25 ºC, 30 min), and digested with endoproteinase LysC (1:10 w:w, 37ºC, over 6h, Wako, cat # 129-02541) and with trypsin (1:10 w:w, 37ºC, over 8h, Promega cat # V5113). After digestion, peptide mixes were acidified with formic acid and desalted with a MicroSpin C18 column (The Nest Group, Inc). A pooled sample was generated by combining equal amounts of the three biological replicates from both conditions to create a pool from which to generate an internal calibration curve with five different serial dilutions. Individual samples were labeled using tandem mass tags (TMT-11) according to the manufacturer instructions and the experimental design specified in Table 1. This experiment was done in triplicate. TMT mixes were fractionated using basic pH reversed-phase fractionation. Twelve fractions were analyzed in using an Orbitrap Eclipse mass spectrometer (Thermo Fisher Scientific, San Jose, CA, USA) coupled to an EASY-nLC 1000 (Thermo Fisher Scientific (Proxeon), Odense, Denmark) with a 90 min gradient.^1^ Data acquisition was done using the real-time synchronous precursos selection MS3 acquisition method (RTS-SPS-MS3).^2^ The scan sequence began with an MS1 spectrum, and in each cycle of data-dependent acquisition analysis, following each survey scan, the most intense ions were selected for fragmentation. Fragment ion spectra were produced via collision-induced dissociation (CID) at normalized collision energy of 35% and they were acquired in the ion trap mass analyzer in “Turbo” mode. MS2 spectra were searched in real time using the algorithm embedded in the instrument control software and the canonical human database from Uniprot (version 2021). MS2 spectra with an Xcorr greater than or equal to 1 and less than 10 ppm precursor mas error, triggered the submission of an MS3 spectrum to the instrument. MS3 spectrum were collected using the multinotch MS3-based TMT method, in a way were ten MS2 fragment ions were captured in the MS3 precursor population using isolation waveforms with multiple frequency notches. MS3 precursors were fragmented by high energy collision-induced dissociation (HCD) at normalized collision energy of 65% and acquired in the Orbitrap analyzer. The mass spectrometry proteomics data have been deposited to the ProteomeXchange Consortium via the PRIDE partner repository with the dataset identifier PXD059628.^3^

Acquired spectra were analyzed using the Proteome Discoverer software suite (v2.4, Thermo Fisher Scientific) and the Mascot search engine (v2.6, Matrix Science).^4^ Data was searched against a customized database including the Uniprot human canonical database plus a list of common contaminants and all the corresponding decoy entries (version 2021).^5^ For peptide identification a precursor ion mass tolerance of 7 ppm was used for MS1 level, trypsin was chosen as enzyme, and up to three missed cleavages were allowed. The fragment ion mass tolerance was set to 0.5 Da for MS2 spectra. Oxidation of methionine and N-terminal protein acetylation were used as variable modifications whereas carbamidomethylation on cysteines, TMT6plex in Lysines and TMT6plex in peptide N-terminal were set as a fixed modification. False discovery rate (FDR) in peptide identification was set to a maximum of 5%. The list of identified peptides was filtered to remove those peptides without quantitation values, i.e. those labelled as “NoQuanLabels”, “NoQuanValues” and “ExcludedByMethod” by the Proteome Discoverer software suite were excluded from the subsequent quantification analysis. Peptides were quantified using the reporter ions intensities in MS3. Reporter ion intensities were adjusted to correct for the isotopic impurities of the different TMT reagents according to manufacturer specifications. A linear regression was fit for peptides with at least three valid quantitative values within the calibration curve. Statistical inference was performed MSstatsTMT (v4.12.1).^6–8^

### Supplementary Figure 1

Venn diagram with the number peptide sequences (with modifications) identified in each TMTCal batch and their overlap.

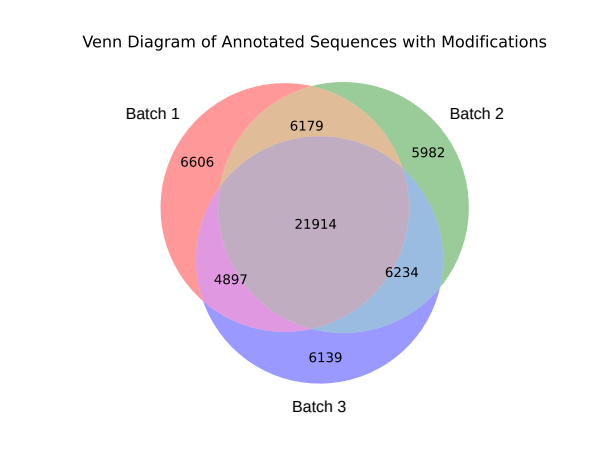

### Supplementary Figure 2

A) Example of the different linear response of peptide [K].TESASVQGR.[N] accross TMTCal batches. The peptide shows five linear quantitative points in two different TMTCal batches and only four in the last TMTCal batch. B) Extracted ion chromatograms for the peptide [K].TESASVQGR.[N] across the three TMTCal batches, demonstrating that the fragmentation event occurred at similar times in the first two TMTCal batches, while in the third batch it was triggered earlier, well before the chromatographic apex. This timing difference may account for the variation in the number of linear quantitative points observed among the batches. C) Boxplots comparing the log-transformed MS3 signal corresponding to the highest concentration in the calibration curves for peptides exhibiting variability in the number of linear quantitative points across batches. The data indicate that, regardless of the precise underlying cause, the signal is consistently lower in the batch where the peptide displays fewer linear quantitative points.

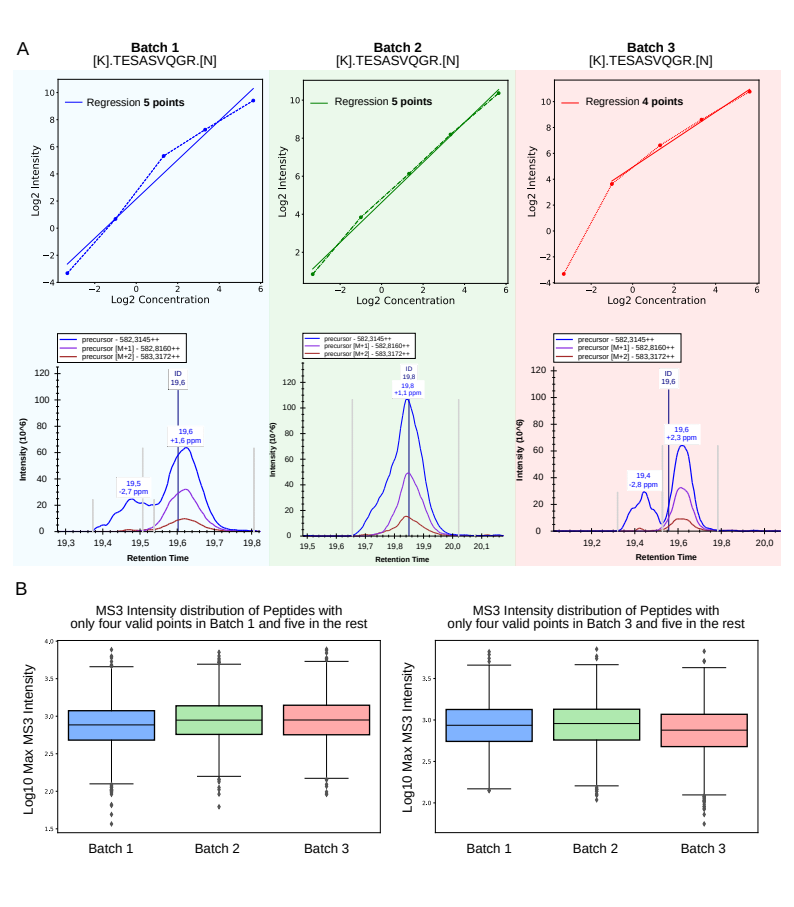

### Supplementary Figure 3

Distribution of the difference (delta) between the logaritmic fold changes obtained with the raw MS3 intensities and those obtained with calibrated MS3 intensities using the TMTCal calibration curve.

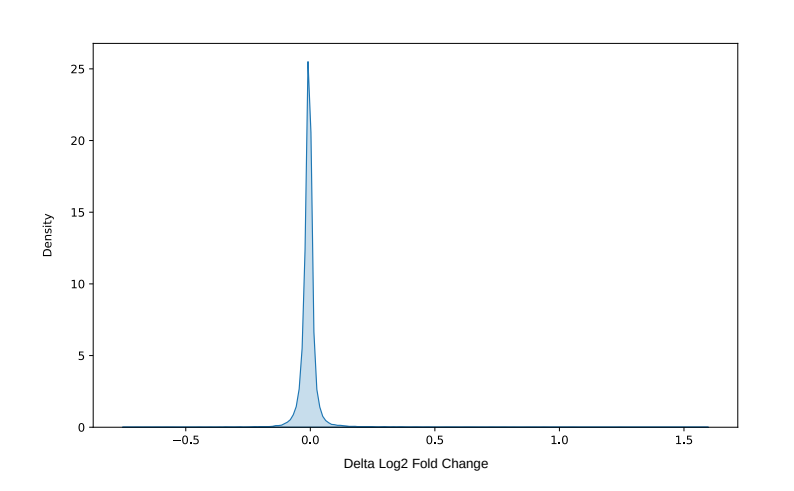

### Supplementary Figure 4

Enrichment plots obtained with the proteins showing a significant change in abundance between untreated and cisplatin-treated (25 µM) SK-OV-3 ovarian cancer cells.

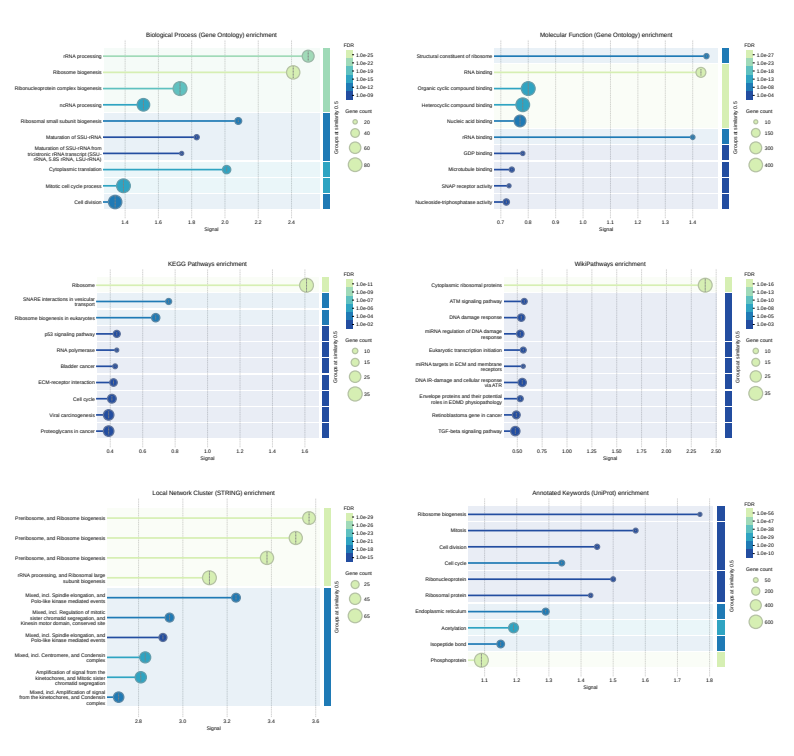

### Supplementary Table 1

List of proteins and peptides identified and quantified in the TMTCal experiments using Proteome Discoverer (v2.4).

### Supplementary Table 2

MSstats relative quantification output with fold-changes and adjusted p-values comparing proteins abundance between untreated and cisplatin-treated (25 µM) SK-OV-3 ovarian cancer cells.
